# Distinct Patterns of Emergence of SARS-CoV-2 Spike Variants including N501Y in Clinical Samples in Columbus Ohio

**DOI:** 10.1101/2021.01.12.426407

**Authors:** Huolin Tu, Matthew R Avenarius, Laura Kubatko, Matthew Hunt, Xiaokang Pan, Peng Ru, Jason Garee, Keelie Thomas, Peter Mohler, Preeti Pancholi, Dan Jones

## Abstract

Following the worldwide emergence of the p.Asp614Gly shift in the Spike (S) gene of SARS-CoV-2, there have been few recurring pathogenic shifts occurring during 2020, as assessed by genomic sequencing. This situation has evolved in the last several months with the emergence of several distinct variants (first identified in the United Kingdom and South Africa) that manifest multiple changes in the S gene, particularly p.Asn501Tyr (N501Y), that likely have clinical impact. We report here the emergence in Columbus, Ohio in December 2020 of two novel SARS-CoV-2 clade 20G variants. One variant, that has become the predominant virus found in nasopharyngeal swabs in the December 2020-January 2021 period, harbors S p.Gln677His (Q677H), affecting a consensus QTQTN domain near the S1/S2 furin cleavage site, nucleocapsid (N) p.Asp377Tyr (D377Y) and membrane glycoprotein (M) p.Ala85Ser (A85S) mutations, with additional S mutations in subsets. The other variant present in two samples, contains S N501Y, which is a marker of the UK-B.1.1.7 (clade 20I/501Y.V1) strain, but lacks all other mutations from that virus. The Ohio variant is from a different clade and shares multiple mutations with the clade 20G viruses circulating in the area prior to December 2020. These two SARS-CoV-2 viruses, which we show are also present and evolving currently in several other parts of North America, add to the diversity of S gene shifts occurring worldwide. These and other shifts in this period of the pandemic support multiple independent acquisition of functionally significant and potentially complementing mutations affecting the S QTQTN site (Q675H or Q677H) and certain receptor binding domain mutations (e.g., E484K and N501Y).

## Introduction

SARS-CoV-2 genomic sequencing has facilitated surveillance efforts to track shifts in viral isolates worldwide (Brufsky, 2020). The emergence in March-April of 2020 of the D614G mutation defining the more transmissible G-strain has been the primary shift during the first nine months of the pandemic (Korber, 2020). This variant has been shown to have increased cell binding and viral spread in *in vitro* models (Mok, 2020; Hu 2020). Within the last several months, however, emergence of several distinct SARS-CoV-2 strains with additional likely pathogenic changes have occurred. These include the rapid spread of novel variants in the United Kingdom (UK, Technical Advisory Group, 2020; European Centre for Disease Prevention and Control, 2020) and South Africa (Tegally, 2020) containing several likely pathogenic but distinct mutations in the Spike (S) gene, particularly N501Y. The rapid transmissibility of these variants (Davies, 2020) and the sudden occurrence of multiple changes in the S gene has raised concerned about shifts in the pattern of COVID-19 disease and possible variability in response to antibody therapies or vaccines.

Here, we report the results of SARS-CoV-2 genomic surveillance from April 2020 through January 2021 in Columbus, Ohio. These data reveals a parallel recent shift in the predominant 20C>20G clade that contains 3 new variants and the emergence of a new virus that harbors the S N501Y variant but is different from the UK-B.1.1.7 (20I/501Y.V1) and the South African variant B.1.351 (20H/501Y.V2). We provide evidence for the geographic localization of both US-based new strains and how they have evolved in late 2020 to early 2021.

## Methods

This study was approved by the Institutional Review Board for the utilization of residual RNA samples from routine clinical SARS-CoV-2 PCR testing for viral sequencing. Briefly, standard PCR-based detection of SARS-CoV-2 was initiated by extraction of viral RNA from nasopharyngeal (NP) swabs (KingFisher™ Flex Magnetic Particle Processor, ThermoFisher). The viral RNA was analyzed, in most cases, using the TaqPath COVID-19 Combo Kit with an Applied Biosystems 7500 Fast Dx Real-Time PCR instrument (ThermoFisher) for SARS-CoV2 detection. SARS-CoV-2 virus sequence was then detected by next-generation sequencing (NGS) using a validated clinical assay in the James Molecular Laboratory at The Ohio State University. Residual RNA from PCR-based testing was reverse-transcribed using SuperScript™ VILO™ cDNA Synthesis Kit (ThermoFisher). NGS was performed using primer sets that tiled the entire SARS-CoV-2 genome (Ion AmpliSeq SARS-CoV-2 Research Panel, ThermoFisher), with library preparation and sequencing performed on Ion Chef and S5, respectively (Ion Torrent, Life Technologies). This panel included primers for the co-amplification of human housekeeping genes to assess RNA quality.

Analysis was performed in the Ion Browser with COVID-19 annotation plugins that produced consensus FASTA files using the IRMA method (reference strain: NC_045512.2). The sequence of the COH.20G/501Y variant in sample DEC32 was confirmed by an independent SARS-CoV-2 genomic sequencing and analysis method. Briefly, RNA was reverse-transcribed using SuperScript™ VILO™ cDNA Synthesis Kit (ThermoFisher). Libraries were produced with KAPA HyperPrep and DI Adapter Kit (Roche). SARS-CoV-2 viral sequences were captured with a COVID-19 Capture Panel covering the entire genome (IDT) and the products were sequenced on the NextSeq 550 (Illumina). The analysis pipeline included BaseSpace, a custom pipeline using GATK tools and DRAGEN RNA Pathogen Detection software (Illumina). Sequences of the two COH.20G/501Y viruses were deposited to GISAID.org as EPI_ISL_832378 and EPI_ISL_826521; an example of COH.20G/677H was deposited as EPI_ISL_826463.

For initial tree-building, individual COVID-19 sequence FASTA files were combined into a single multifasta file with a custom shell script. The multifasta files were aligned using MAFFT (Katoh, 2002) (version 7.453) using default settings. MAFFT alignment files were analyzed for maximum likelihood using RAxML (Stamatakis, 2006) (version 8.2.12) using the GTRGAMMA model with 1000 bootstraps. The tree was produced using Dendroscope (Huson and Scornavacca 2012) (version 3.7.2) with default settings. Numbers at the tree branches in Supplementary Figure 1 represent percent of bootstraps supporting a branch (i.e. 30 = 300/1000 runs supporting this branch). Strain typing and clades were designated using the most recent NextStrain nomenclature, with clade designation as 20G throughout if the clade-defining mutations listed in Supplementary Table 1 were present (Bedford, 2021).

For extended phylogenetic analysis (Figures 1 and 2, and supplementary Figures 2 and 3), two large datasets were generated using aligned data provided by GISAID: Q667H (356 sequences, downloaded on 1/22/21) and N501Y (132 sequences, downloaded on 1/23/21). Prior to formal phylogenetic analyses, both data sets were examined in TempEst version 1.5.3 (Rambaut, 2016) to screen for outliers and assess temporal signal. Phylodynamic analyses were then carried out in BEAST version 1.10.4 (Suchard, 2018) under a strict clock model with coalescent exponential growth and an HKY substitution model with gamma-distributed rates (4 categories) and unlinked parameters for each codon position. The prior distribution for the exponential growth rate parameter was set to Laplace(0,100) and the prior for the exponential population size parameter was set to gamma(0,10). Default priors were used for all other parameters. The Q677H dataset was run for 40 million iterations, with the first 4 million discarded as burnin and every 1,000^th^ tree retained. The N501Y dataset was run for 20 million iterations, with the first 2 million discarded as burnin and every 1,000^th^ tree retained. Tracer version 1.7.1 (Rambaut, 2018) was used to assess convergence. For the Q677H dataset, ESS values were above 200 for all but four parameters, with all four of these having values above 150. The ESS values for all parameters in the N501Y analysis were above 200. TreeAnnotater version v1.10.4 (Suchard, 2018) was used to compute the maximum clade credibility (MCC) tree, which was plotted using FigTree version 1.4.4 (Rambaut, 2018).

**Figure 1.**
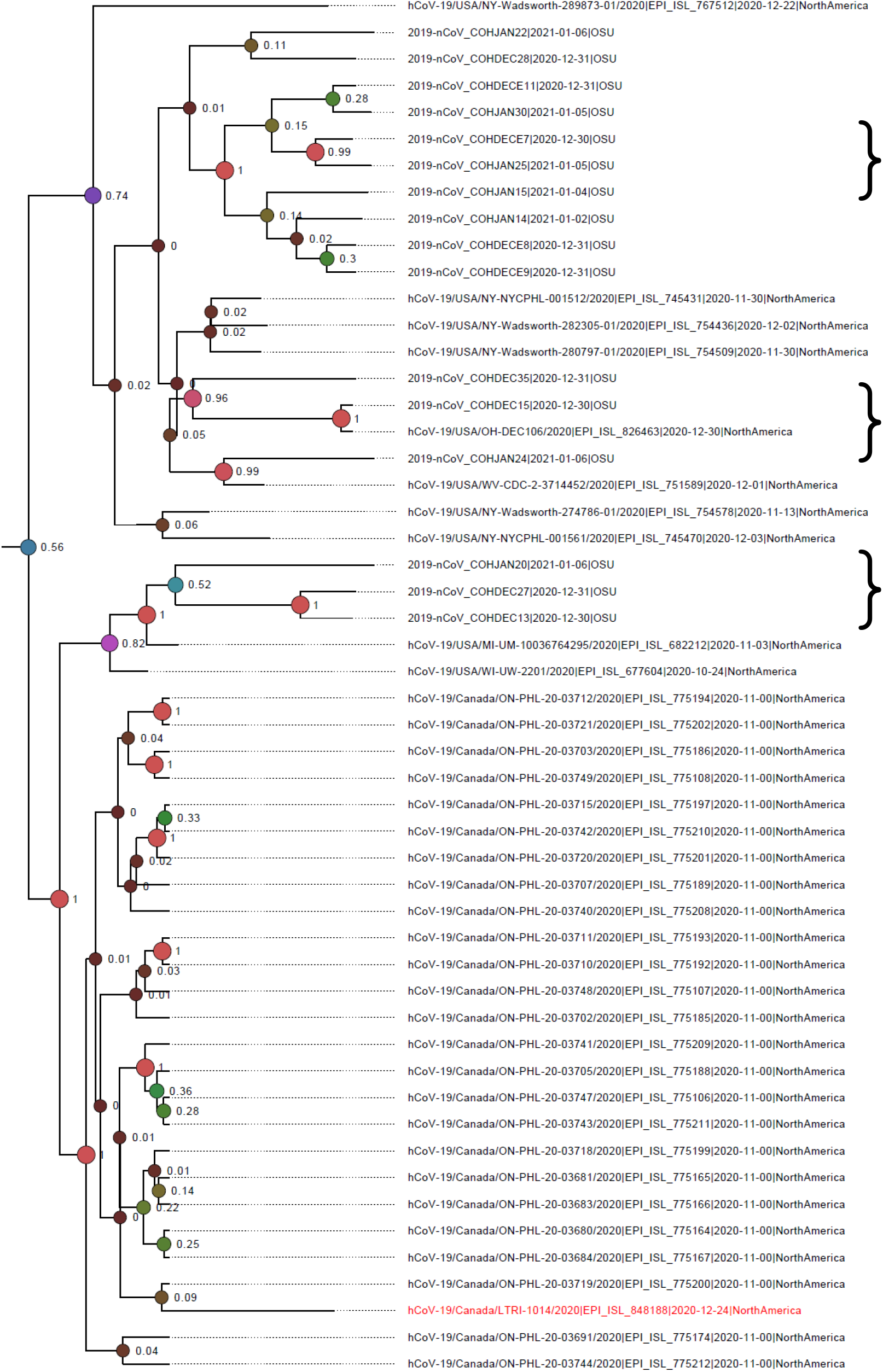
Phylogenic relationship between the 20G/677H viruses seen in Columbus and other GISAID-reported clade GH viruses over time. This figure represents the 20G clade section of a more extensive phylogenetic analysis of Q677H-bearing viruses reported worldwide in GISAID as of January 22, 2021. The full tree is provided in radial form in Supplementary Figure 2. Brackets indicates a representative sampling of 20G/677H samples from Columbus, Ohio. The case in red ink is a highly similar virus collected in Toronto, Canada on December 24, 2020 (EPI_ISL_848188) that was reported to also show S E484K. The color and width of the circles at the internal nodes indicate posterior probability, with “warmer colors” and larger circles indicating higher support. Red circles indicate posterior probabilities of 1.

**Figure 2.**
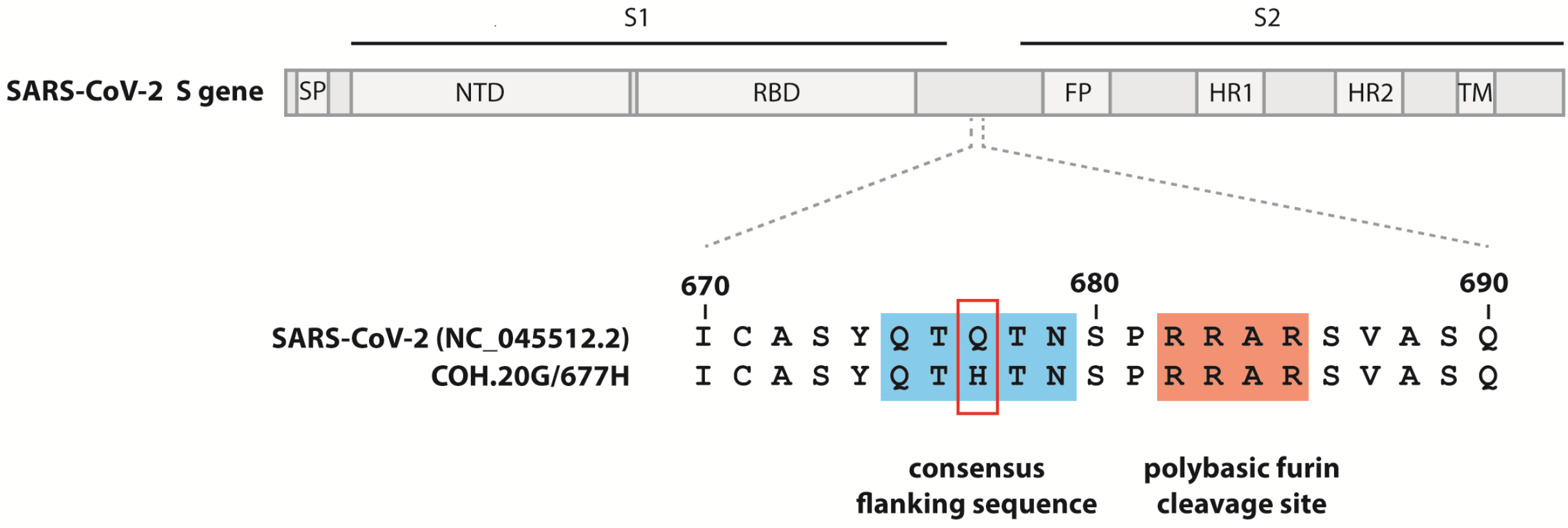
Site of the Q677H mutation in Spike gene in the QTQTN motif conservation/furin cleavage site. Signal peptide (SP), N-terminal domain (NTD), receptor-binding domain (RBD), fusion peptide (FP), heptad repeat 1 (HR1), heptad repeat 2 (HR2), and transmembrane domain (TM).

## Results

### Summary of sequencing results in the early and mid-pandemic period

RNA extracted from PCR-positive nasopharyngeal samples from the Columbus OH area was sequenced in April (n= 56), May (n = 71), June (n=21), July (n=16), September (n = 11) and December 2020 (n =36) and January 2021 (n = 24) for surveillance purposes. A total of 235 NP samples were sequenced. In April 2020, two samples were positive for the S-strain with the remainder representing the G strain.

Aside from the G strain-defining changes (Supplementary Table 1), there were very few recurrent non-conservative/non-synonymous changes observed in April and May 2020 samples. In that period, most of the G-strain positive cases represented G strain alone or an unspecified G branch (17.3%, commonly with ORF8 p.Ala51Val) or the 20C clade bearing ORF3 p.Gln57His (80.3%), with few representing clade 20B (2.4%). In June and July 2020, as apparent infection rates in Columbus decreased, there was a proportional increase in clade 20B (40.5% of samples), with fewer clade G/unspecified (21.7%) and clade C viruses (37.8%). In September, coinciding with an increase in PCR positivity rates in the area, 20C clade samples again predominated (72.7% of samples), with some showing additional variants closely matching the newly designated NextStrain 20G clade (Bedford, 2021), with the remaining being 20A or 20B clade viruses. Samples were not obtained during the months of October and November 2020.

### Rapid emergence of a clade 20G virus with shared S, N and M mutations

When sequencing resumed in late December, we noted the emergence of a distinct 20G clade that had acquired the following variants: S p.Gln677His (Q677H), M p.Ala85Ser (A85S) and N p.Asp377Tyr (D377Y, Table 1A) and is designated COH.20G/677H (Figure 1, brackets). During the week of Dec 21^st^ 2020, these 3 variants was co-detected in 1 of 10 samples (10%), but were detected in 6/20 (30%), 6/10 (60%) and 8/13 (61.5%) of samples in the following weeks. Three virus samples also showed N D377Y without the other two changes, indicating it was an earlier change (Supplementary Figure 1). Other changes seen in a subsets of COH.20G/677H cases included ORF1AB p.Ile529Val and S p.Thr95Ile followed by either S p.Leu5Phe or S p.Asn914Ser.

**Table 1A.** Mutations present in the emergent clade 20G/677H SARS-CoV-2 virus from Columbus Ohio samples.

| Gene | Nucleotide | cDNA | Amino acid | Cases | % in last month |
| --- | --- | --- | --- | --- | --- |
| N | G29402T | c.1129G>T | D377Y | 24 | 38.1 |
| S | G23593T | c.2031G>T | Q677H | 21 | 33.3 |
| M | G26775T | c.253G>T | A85S | 19 | 30.2 |
| **Subgroup** |  |  |  |  |  |
| ORF1AB | A1850G | c.1585A>G | I529V | 8 | 12.7 |
| S | C21846T | c.284C>T | T95I | 10 | 15.9 |
| **Minor (1)** |  |  |  |  |  |
| S | C21575T | c.13C>T | L5F | 3 | 4.8 |
| **Minor (2)** |  |  |  |  |  |
| S | A25563G | c.2741A>G | N914S | 3 | 4.8 |

In all cases, these co-occurring variants arose in a 20G clade variant branch that had been present in Columbus since at least September 2020. The backbone was defined by ORF1AB: p.Met2606Ile (c.7818G>A), p.Leu3352Phe (c.10054C>T), p.Thr4847Ile (c.14540C>T), p.Leu6053Leu (c.18159A>G), p.His7013His (c.21039C>T); ORF3A: p.Gly172Val (c.515G>T); ORF8: p.Ser24Leu (c.71C>T); N: p.Pro67Ser (c.199C>T) and p.Pro199Leu (c.596C>T).

Phylogenetic analysis of S Q677H-bearing viruses present in GISAID, as of January 23 2021, shows a large number of similar virus (>250 instances) have now been reported from 11 different states in late 2020 and January 2021 (Figure 1 and Supplementary Figure 2), with the Columbus samples most similar to samples reported from Ontario, Canada and upstate New York, which are geographically contiguous areas to Ohio.

### Emergence of a distinctive clade 20G virus harboring S N501Y

In late December (12/30/20) and January (1/6/21), we detected two samples with a 20G strain backbone that had acquired S N501Y, as well as ORF8 R52I in one case (designated COH.20G/501Y, Figure 1 arrow and Figure 3, red ink) which are both changes present in the UK-B.1.1.7 strain. In contrast to the B.1.1.7 strain, which has a 20B origin, the S N501Y and ORF8 R52I variants identified in this case were on a 20G background common to our area, as defined by ORF1AB: p.Leu3352Phe (c.10054C>T), p.Leu6053Leu (c.18159A>G), p.His7013His (c.21039C>T), ORF3A: p.Gly172Val (c.515G>T), ORF8: p.Ser24Leu (c.71C>T), N: p.Pro67Ser (c.199C>T), and p.Pro199Leu (c.596C>T). Both samples also shared mutations in ORF1AB (p.Thr999Ile, p.Ala1074Val and p.Leu1313Leu, Table 1B) indicating they likely arose from the same precursor. In addition to lacking the characteristic N p.ArgGly203LysArg (c.608_610delGGGinsAAC) marking the 20B clade, COH.20G/501Y viruses also lack the other mutations seen in B.1.1.7, as summarized in Supplementary Table 1.

**Figure 3.**
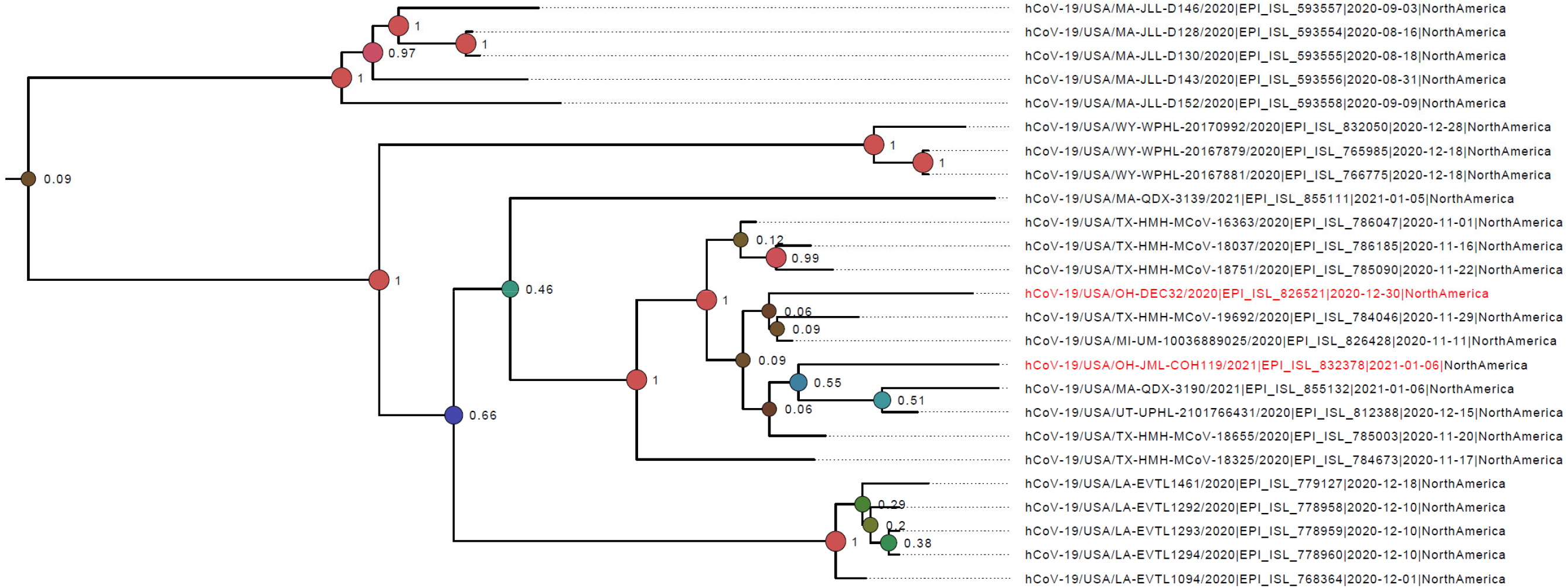
Phylogenic relationship between the 20G/501Y viruses seen in Columbus and across the United States. This figure represents the 20G clade section of an extended phylogenetic tree displaying all N501Y-bearing strains reported in GISAID in the United States as of January 23, 2021. The full tree, including mostly B.1.1.7 examples, is included in Supplementary Figure 2. Samples with red ink are nasopharyngeal swabs from patients tested in Columbus, Ohio on 12/30/20 and 1/6/21. Adjacent samples are similar 20G viruses with S 501Y but no other recurrent S RBP mutations from six other states and had collection dates in late 2020 and early 2021. See Methods for details on tree-building and interpretation. Color and width of circles are as in Figure 1.

**Figure 4.**
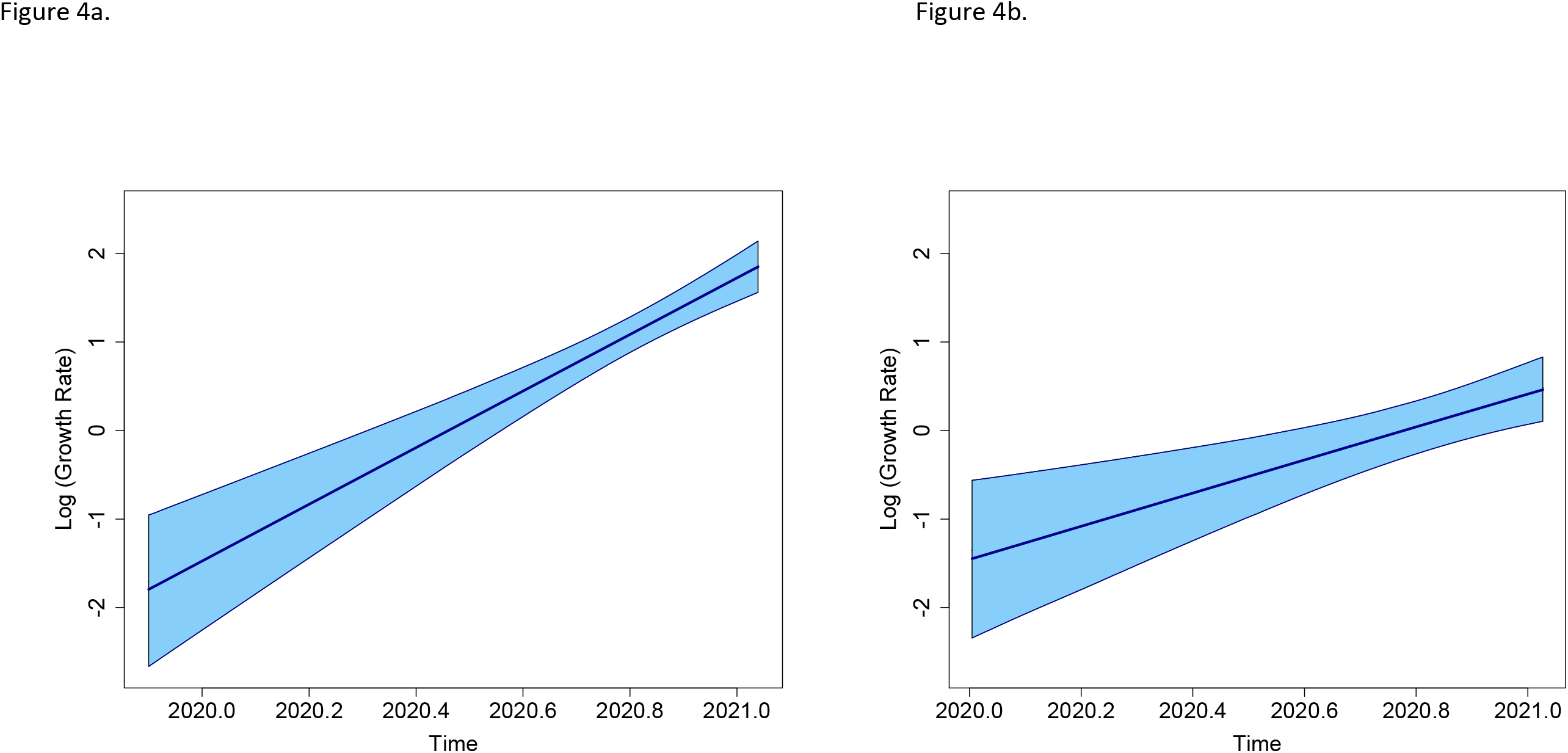
Exponential growth rate estimates of SARS-CoV-2 viruses in Ohio suggest increasing transmission rates. Median (center line) of the estimated growth rate parameter in the exponential model from the phylodynamic analyses with 95% HPD intervals (shaded) for the Q677H dataset (a) and the N501Y dataset (b). The increasing trend through time is indicative of the recent increase in the rate of transmission of these viruses.

**Table 1B.** Novel variants in the 20G/501Y sample in Columbus and other sites in the United States. Includes common changes in COH.20G/501Y (EPI_ISL_832378 and EPI_ISL_826521) and similar viruses reported by other groups, including EPI_ISL_826428, EPI_ISL_812388, EPI_ISL_784046 (See Discussion for details).

| Gene | nucleotide | cDNA | amino acid |
| --- | --- | --- | --- |
| ORF1AB | C3261T | c.2996C>T | T999I |
|  | C3486T | c.3221C>T | A1074V |
|  | C4202T | c.3937C>T | L1313L |
| S | A23063T | c.1501A>T | N501Y |
| **Present in 1 of 2 viruses** |  |  |  |
| ORF8 | G28048T | c.155G>T | R52I |

Phylogenetic analysis of viruses present in GISAID as of January 23 2021 shows multiple sequences that are highly similar to COH.20G/501Y are now reported from 6 different states in late 2020 and January 2021 (Figure 3 and Supplementary Figure 3). These represent the only 20G-S 501Y-bearing viruses reported in United States except for a small cluster of distinct viruses from Massachusetts collected in August 2020 that have not been reported since that time (e.g. EPI_ISL_593557, top branch in Figure 3). As discussed above, extended phylogenetic analysis shows that the N501Y-containing B.1.1.7 viruses now present in the United States are completely distinct (Supplementary Figure 3).

## Discussion

We report two samples in which SARS-Co2 had acquired the S N501Y variant in late December 2020/January 2021 in Columbus Ohio. This particular amino acid change was first highlighted in a clinical sample in the United Kingdom in association with other novel S variants and a clade 20B backbone (ECDC, 2020; Davies, 2020), with the combination named as the B.1.1.7 strain and a Next Strain designation as 20I/501Y.V1 (Bedford, 2021). The same N501Y mutation was subsequently found in a clade 20C strain in South Africa, where it was associated with a different set of additional S variants (Tegally, 2020), with Next Strain designation as 20H/501Y.V2 or B.1.351. In late December 2020, the incidence of detection of both of these variants began markedly increasing, implicating S 501Y (with or without other S mutations) in increased transmissibility.

The two Columbus viruses with S N501Y (COH.20G/501Y) have a 20G backbone but lack nearly all of the reported consensus changes in 20I/501Y.V1 as well as those in the 20H/501Y.V2. They cluster away from other S501Y-bearing strains in an extended phylogenetic analysis (Supplementary Figure 2). This favors an independent acquisition of this variant in a 20G clade branch that has been consistently present in Ohio since at least September 2020. Emergence of these two new variants was correlated with months when the diversity of viruses reported in Ohio in the GISAID database was increasing (Figure 3).

Since the first version of this pre-print, nearly identical N501Y-bearing viruses (all currently classified as B.1.2/GH by NextStrain) have now been reported in Massachusetts (multiple submissions, including EPI_ISL_855132), Michigan (EPI_ISL_826428), Utah (EPI_ISL_812388) and Texas (multiple submissions, including EPI_ISL_784046), as of January 23, 2021. Related viruses with additional mutations were also reported from Wyoming (e.g., EPI_ISL_765985) and Louisiana (EPI_ISL_778960). These viruses have collection dates from early November 2020 to mid-January 2021 compatible with spread of this variant throughout the United State during that period. These represent the only 20G/501Y-bearing viruses reported in United States as of January 23, 2021, except for a small cluster of distinct viruses from Massachusetts from August 2020 that have not been seen since that time (e.g. EPI_ISL_593557, Figure 3 top).

The S N501Y mutation, located within the receptor binding domain, is of particular concern for two reasons. First, the S protein with 501Y mutation displays increased affinity for ACE2 (Luan, 2020; Starr, 2020). Second, S 501Y mutation may impact association of receptor binding neutralizing antibodies including those in the Regeneron cocktail (Weisblum, 2020; Starr, 2020). The S N501Y mutation has also been shown to emerge spontaneously with viral passaging in a mouse model of SARS-CoV-2 infection (Gu, 2020), supporting its role in promoting viral spread and/or transmissibility. The only other shared mutation of COH.20G/501Y with B.1.1.7, in one case only, is the ORF8 R52I mutation. B.1.1.7 also has a deletion involving ORF8 that would likely inactivate its functions; such deletions have emerged in multiple strains of SARS-CoV-2 (Su, 2020) but were not present in COH.20G/501Y.

We also report the emergence of a predominant SARS-CoV-2 strain with a 20G clade backbone that has single mutations in the S, M and N genes (Q677H, A85S and D377Y, respectively), first detected in Columbus, Ohio in December 2020. Since the first publication of this pre-print, there have been reports of closely related 20G variants also harboring S Q677H extensively present in several states in the upper Midwest, including Michigan, Wisconsin and Minnesota (Pater, 2021) and Ontario, Canada. There are over 250 highly-related viruses reported now in GISAID as of January 23, 2021. One of these nearly identical 20G/677H viruses reported from Toronto (EPI_ISL_848188) has acquired the S Glu484Lys (E484K) mutation located in the RBP (red highlight in Figure 1/Supplementary Figure 2). This mutation which is a key change in the unrelated 20H/501Y.V3 strain has been shown to increase binding of S to human ACE2 *in vitro* (Nelson, 2021) and to interfere with binding of some but not all natural-occurring and therapeutic antibodies directed against the Spike RBP domain (Hu, 2021; Greaney, 2021; Shi 2021).

The S Q677H mutation defining this emergent strain disrupts a QTQTN consensus sequence (Figure 2) adjacent to the polybasic furin cleavage site spanning the S1 and S2 junction (Jacob, 2020). Prior to late 2020, that mutation has been reported in NextStrain as a sporadic variant arising in multiple strains (Kim, 2020). Since late 2020, the S Q677H mutation has now also been reported in over 80 20I/501Y.V1 viruses (S N501Y-bearing) in GISAID as of January 23, 2021 (See branch in Supplementary Figure 2). This suggests that these two different types of S mutations (RBP and QTQRTN motif) may have complementary actions on virus function.

The other non-conservative mutation altering the Spike QTQTN motif currently prevalent is S Q675H, which has been detected in over 1200 viruses from GISAID as of January 23, 2021, and is a marker of the B.1.243/G strain emerging on the West Coast of the US. Dual Q675H and Q677H have also been reported in 35 B.1.36/GH viruses since November 2020 and only very rarely previously. Deletions spanning the QTQTN motif have also been reported and may influence viral properties (Liu, 2020).

Among the other shared mutations in the consensus 20G/677H virus backbone (Table 1A), N D377Y is the most commonly reported (Gupta, 2020) and is a marker of the unrelated European-based B.1.177/GV, B.1.236/G and B.1.258/G clades. M A85S is an uncommon mutation, largely seen to date only in the 20G/677H strain and in the unrelated European-based B.1.236/G clade. Based on analysis of viruses from Columbus and other states, N D377Y appears to have preceded S Q677H followed by M A85S in the outgrowth of this strain. The rapid emergence in the last 2 months of the 20G/677H strain across the Midwestern United States and Canada, now with at least one instance of acquisition of E484K, merits close attention.

## Supporting information

FASTA for 20G/677H example

FASTA for 20G/501Y example

## References

Bedford T, Hodcroft EB, Neher RA, Artic Network. Updated Nexstrain SARS-CoV-2 clade naming strategy. Available at https://virological.org/t/updated-nextstain-sars-cov-2-clade-naming-strategy/581, accessed 2021 Jan 11.

Brufsky A. Distinct Viral Clades of SARS-CoV-2: Implications for Modeling of Viral Spread. Journal of medical virology. 2020 Apr 20.

Davies NG, Barnard RC, Jarvis CI, Kucharski AJ, Munday J, Pearson CA, Russell TW, Tully DC, Abbott S, Gimma A, Waites W. Estimated transmissibility and severity of novel SARS-CoV-2 Variant of Concern 202012/01 in England. medRxiv. 2020 Dec 26.

ECDC: European Centre for Disease Prevention and Control. Rapid increase of a SARS-CoV-2 variant with multiple spike protein mutations observed in the United Kingdom – 20 December 2020. ECDC: Stockholm; 2020.

Greaney AJ, Loes AN, Crawford KH, Starr TN, Malone KD, Chu HY, Bloom JD. Comprehensive mapping of mutations to the SARS-CoV-2 receptor-binding domain that affect recognition by polyclonal human serum antibodies. bioRxiv. 2021 Jan 4.

Greaney AJ, Starr TN, Gilchuk P, Zost SJ, Binshtein E, Loes AN, Hilton SK, Huddleston J, Eguia R, Crowe Jr JE, Bloom JD. Complete mapping of mutations to the SARS-CoV-2 spike receptor-binding domain that escape antibody recognition. bioRxiv. 2020 Sept 10.

Gu H, Chen Q, Yang G, He L, Fan H, Deng YQ, Wang Y, Teng Y, Zhao Z, Cui Y, Li Y. Adaptation of SARS-CoV-2 in BALB/c mice for testing vaccine efficacy. Science. 2020 Sep 25;369(6511):1603–7.

Gupta A, Sabarinathan R, Bala P, Donipadi V, Vashisht D, Katika MR, Kandakatla M, Mitra D, Dalal A, Bashyam MD. Mutational landscape and dominant lineages in the SARS-CoV-2 infections in the state of Telangana, India. medRxiv. 2020 Aug 26.

Hu J, He CL, Gao Q, Zhang GJ, Cao XX, Long QX, Deng HJ, Huang LY, Chen J, Wang K, Tang N. The D614G mutation of SARS-CoV-2 spike protein enhances viral infectivity. bioRxiv. 2020 Jan 1.

Hu J, Peng P, W K, Liu B-z, Fang L, Luo F0y, Jin A-s, Tang N, Huang A. Emerging SARS-CoV-2 variants reduce neutralization sensitivity to convalescent sera and monoclonal antibodies. bioRxiv. 2021 Jan 22.

Huson DH, Scornavacca C. Dendroscope 3: an interactive tool for rooted phylogenetic trees and networks. Systematic biology. 2012 Dec 1;61(6):1061–7.

Jacob JJ, Vasudevan K, Pragasam AK, Gunasekaran K, Kang G, Veeraraghavan B, Mutreja A. Evolutionary tracking of SARS-CoV-2 genetic variants highlights intricate balance of stabilizing and destabilizing mutations. bioRxiv. 2020 Dec 29.

Katoh K, Misawa K, Kuma KI, Miyata T. MAFFT: a novel method for rapid multiple sequence alignment based on fast Fourier transform. Nucleic acids research. 2002 Jul 15;30(14):3059–66.

Kim JS, Jang JH, Kim JM, Chung YS, Yoo CK, Han MG. Genome-Wide Identification and Characterization of Point Mutations in the SARS-CoV-2 Genome. Osong Public Health and Research Perspectives. 2020 Jun;11(3):101.

Korber B, Fischer WM, Gnanakaran S, Yoon H, Theiler J, Abfalterer W, Hengartner N, Giorgi EE, Bhattacharya T, Foley B, Hastie KM. Tracking changes in SARS-CoV-2 Spike: evidence that D614G increases infectivity of the COVID-19 virus. Cell. 2020 Aug 20;182(4):812–27.

Liu Z, Yuan R, Li M, Lin H, Peng J, Xiong Q, Sun J, Li B, Wu J, Hulswit RJ, Bowden TA. Identification of a common deletion in the spike protein of SARS-CoV-2. J Virol. 22 June 2020.

Luan, B., Wang, H. and Huynh, T., Molecular Mechanism of the N501Y Mutation for Enhanced Binding between SARS-CoV-2’s Spike Protein and Human ACE2 Receptor. bioRxiv. 2021 Jan 5.

Mok BW, Cremin CJ, Lau SY, Deng S, Chen P, Zhang AJ, Lee AC, Liu H, Liu S, Ng TT, Lao HY. SARS-CoV-2 spike D614G variant exhibits highly efficient replication and transmission in hamsters. bioRxiv. 2020 Aug 28. Nelson G, Buzko O, Spilman PR, Niazi K, Rabizadeh S, Soon-Shiong PR. Molecular dynamic simulation reveals E484K mutation enhances spike RBD-ACE2 affinity and the combination of E484K, K417N and N501Y mutations (501Y. V2 variant) induces conformational change greater than N501Y mutant alone, potentially resulting in an escape mutant. bioRxiv. 2021-01.

Pater AA, Bosmeny MS, Barkau CL, Ovington KN, Chilamkurthy R, Parasrampuria, Eddington SB, Yinusa AB, White AA, Metz PE, Sylvain RJ, Hebert MM, Benzinger SW, Sinha K, Gagnon KT. Emergence and evolution of a prevalent new SARS-CoV-2 variant in the United States. bioRxiv. 2021 Jan 13.

Rambaut A, Lam T, Carvallo LM, and Pybus OG. Exploring the temporal structure of sequences using TempEst, Virus Evolution 2(1): vew007, 2016.

Rambaut A. FigTree, available at https://github.com/rambaut/figtree, 2018.

Rambaut A, Drummond AJ, Xie D, Baele G and Suchard MA (2018) Posterior summarisation in Bayesian phylogenetics using Tracer 1.7. Systematic Biology 67(5): 901–904, 2018.

Santos JC, Passos GA. The high infectivity of SARS-CoV-2 B. 1.1. 7 is associated with increased interaction force between Spike-ACE2 caused by the viral N501Y mutation. bioRxiv. 2021 Jan 1:2020–12.

Shi PY, Xie X, Zou J, Fontes-Garfias C, Xia H, Swanson K, Cutler M, Cooper D, Menachery V, Weaver S, Dormitzer P. Neutralization of N501Y mutant SARS-CoV-2 by BNT162b2 vaccine-elicited sera Neutralization of N501Y mutant SARS-CoV-2 by BNT162b2 vaccine-elicited sera. 2021 Jan 7

Stamatakis A. RAxML-VI-HPC: maximum likelihood-based phylogenetic analyses with thousands of taxa and mixed models. Bioinformatics. 2006 Nov 1;22(21):2688–90.

Starr TN, Greaney AJ, Hilton SK, Ellis D, Crawford KH, Dingens AS, Navarro MJ, Bowen JE, Tortorici MA, Walls AC, King NP. Deep mutational scanning of SARS-CoV-2 receptor binding domain reveals constraints on folding and ACE2 binding. Cell. 2020 Sep 3;182(5):1295–310.

Su Y, Anderson D, Young B, Zhu F, Linster M, Kalimuddin S, Low J, Yan Z, Jayakumar J, Sun L, Yan G. Discovery of a 382-nt deletion during the early evolution of SARS-CoV-2. bioRxiv. 2020 Mar 12.

Suchard MA, Lemey P, Baele G, Ayres DL, Drummond AJ & Rambaut A. Bayesian phylogenetic and phylodynamic data integration using BEAST 1.10 Virus Evolution 4, vey016, 2018.

Technical Advisory Group: Brief on the viral variant VOC-202012/01, 23 December 2020.

Tegally H, Wilkinson E, Giovanetti M, Iranzadeh A, Fonseca V, Giandhari J, Doolabh D, Pillay S, San EJ, Msomi N, Mlisana K. Emergence and rapid spread of a new severe acute respiratory syndrome-related coronavirus 2 (SARS-CoV-2) lineage with multiple spike mutations in South Africa. medRxiv. 2020 Dec 22.

Weisblum Y, Schmidt F, Zhang F, DaSilva J, Poston D, Lorenzi JC, Muecksch F, Rutkowska M, Hoffmann HH, Michailidis E, Gaebler C. Escape from neutralizing antibodies by SARS-CoV-2 spike protein variants. eLife. 2020 Oct 28;9:e61312.

