## Supplementary Figure 1 for "Distinct Patterns of Emergence of SARS-CoV-2 Spike Variants including N501Y in Clinical Samples in Columbus Ohio"

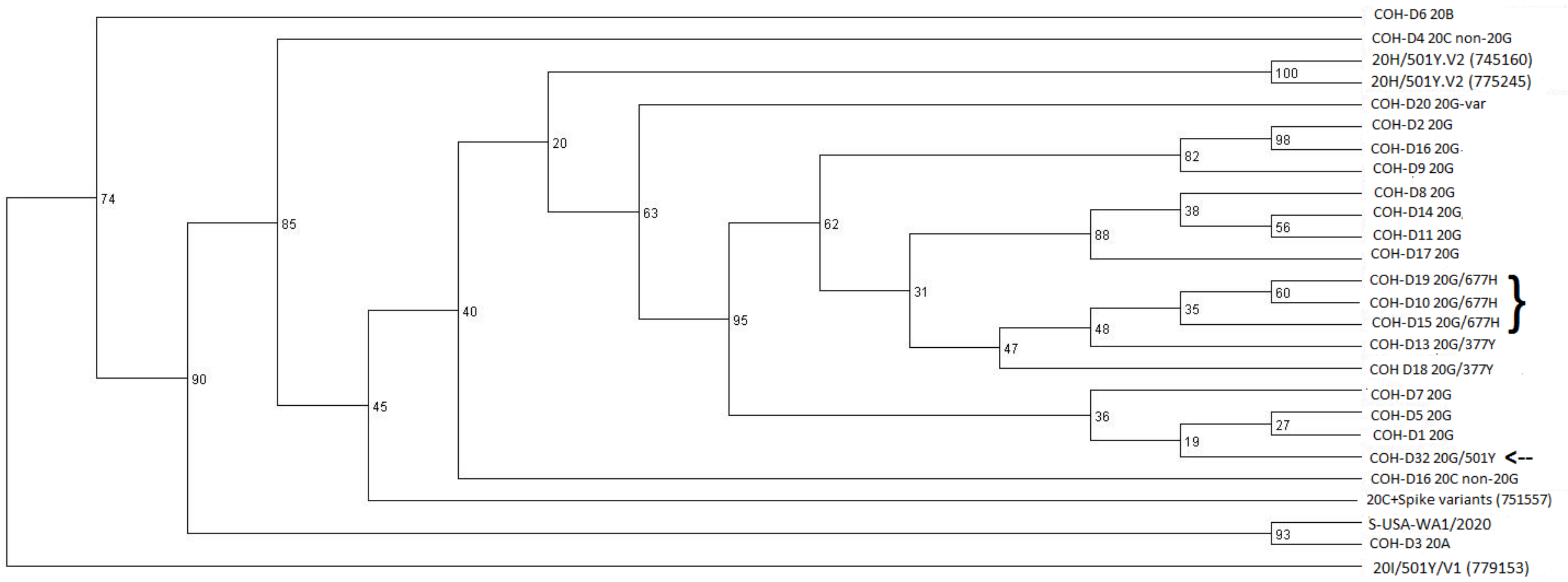

**Supplementary Figure 1. Phylogenic relationship between viruses seen in late December in Columbus Ohio.** Samples labeled "COH" are nasopharyngeal swabs from patients tested in Columbus, Ohio from 12/21-12/31/20. Most are clade 20G, with one each being 20A, 20B, and a 20G variant (var) that does not show every strain-defining mutation. One of the 20G/501Y viruses is included and marked with an arrow. Three examples of the emerging 20G/677H variant are bracketed, with the adjacent 20G/377Y viruses containing N D377Y but not S Q677H or M A85S. Reference sequences (FASTA files downloaded from GISAID.org) show recent examples of South Africa B.1.351/20H/501Y.V2 (EPI_ISL_745160, 12/4/2020), a B.1.351 20H/501Y.V2 strain collected in Australia (EPI_ISL_775245, 1/4/2021), a B.1.1.7/20I/501Y.V1 virus collected in the United States (EPI_ISL_779154, 1/4/2021) and a 20C-derived virus from Nevada with several distinct S variants (EPI_ISL_751557, 12/4/2020). The *2019-nCoV*/USA-WA1/2020 SARS-CoV-2 reference stain (ATCC) was used as a sequencing control. See Methods for details on tree-building and interpretation.
