## Supplementary Figure 2 for "Distinct Patterns of Emergence of SARS-CoV-2 Spike Variants including N501Y in Clinical Samples in Columbus Ohio"

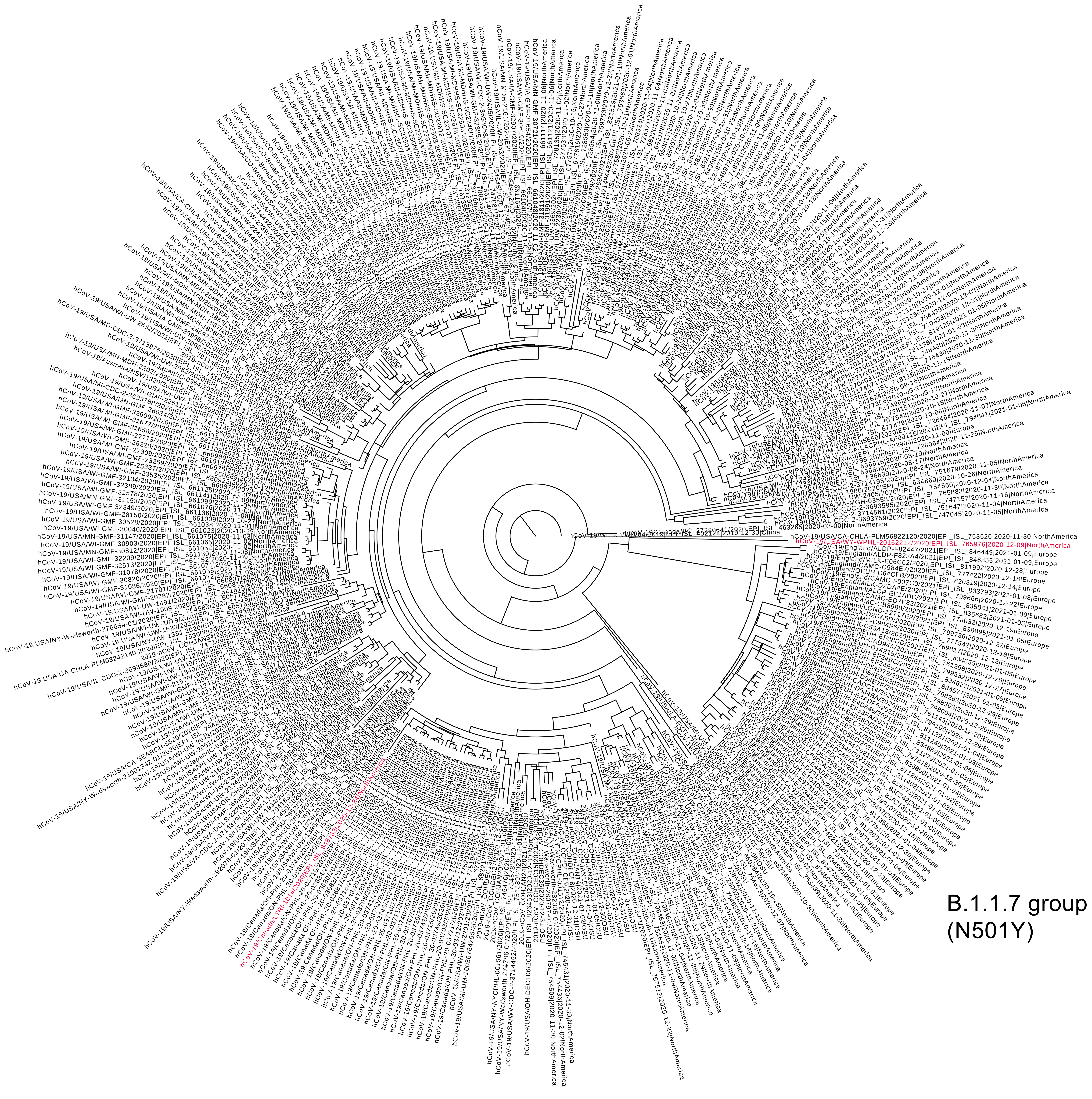


Clade 20G (area in Figure 1)

**Supplementary Figure 2. Extended analysis of SARS-CoV-2 viruses containing S Q677H mutation worldwide as of January 23, 2021.** The major radial branch is the 20G/677H strain centered in the Midwest United States, with the area show in Figure 1 marked at the bottom. The area major S Q677H-bearing group is B.1.1.7/20I/501Y.V1 viruses (right). Red ink highlights a 20G/677H virus that also bears S E484K and another virus reported from Wyoming that also showed S E484K (EPI_ISL_765976). Sequences representing the Wuhan L strain (EPI_ISL_402124) and G strain clades O (EPI_ISL_753526), G (EPI_ISL_463265 and EPI_ISL_583293) and GV (EPI_ISL_649919) that all do not contain Q677H are included for reference. See Methods for details on case selection, tree-building and interpretation.
