## Supplementary Figure 3 for "Distinct Patterns of Emergence of SARS-CoV-2 Spike Variants including N501Y in Clinical Samples in Columbus Ohio"

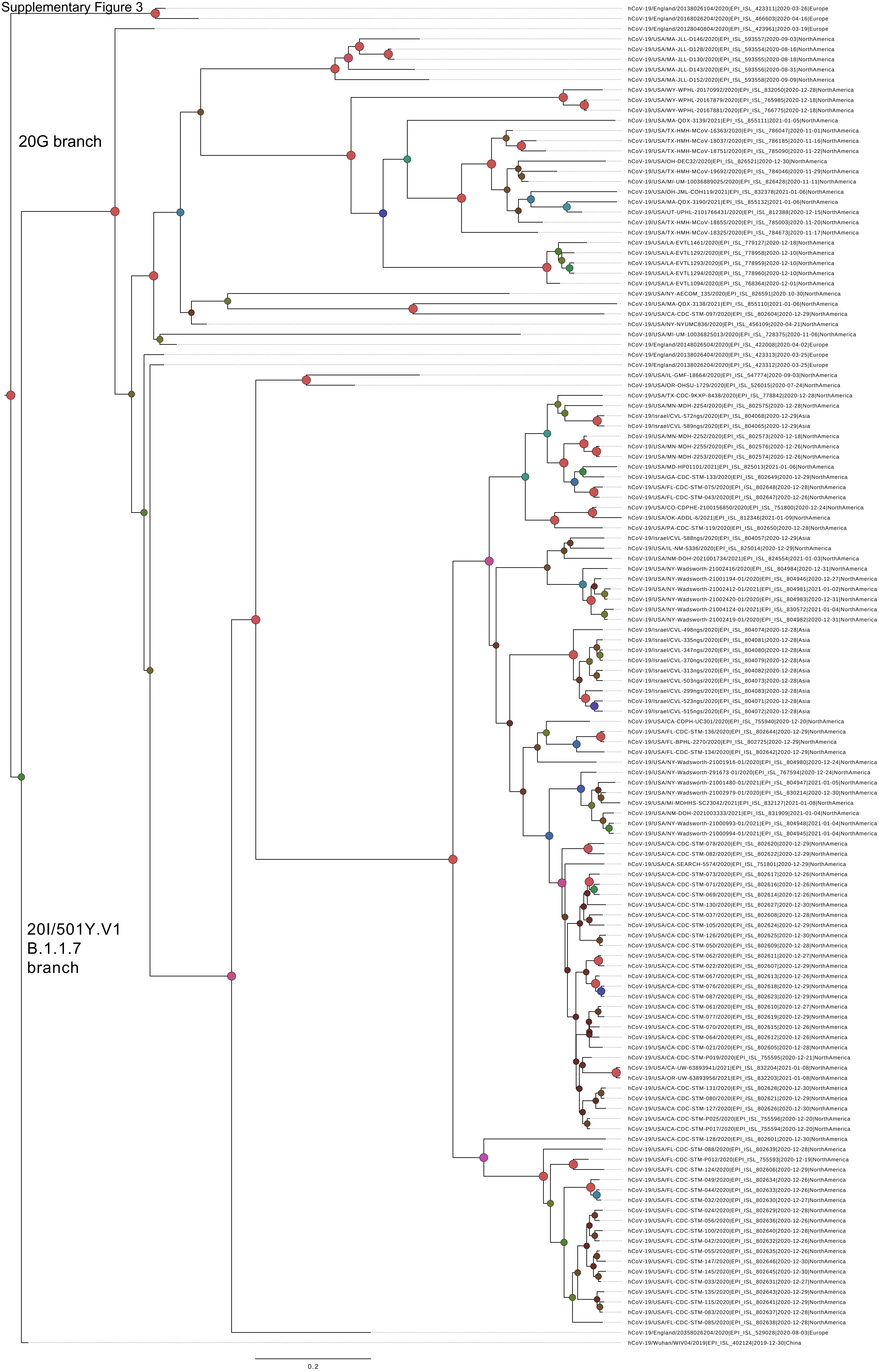

**Supplementary Figure 3. Extended analysis of SARS-CoV-2 viruses containing S N501Y in the United States as of January 23, 2021.** The top branches are 20G clade viruses, whereas the bottom branches are B.1.1.7/20I/501Y.V1 strain. Red ink highlights the 20G/501Y viruses from Columbus, Ohio. See Methods for details on tree-building and interpretation. Sequences representing the Wuhan L-strain (EPI_ISL_402124), G-strain clades O (EPI_ISL_466603), V (EPI_ISL_423311), G (EPI_ISL_423961, EPI_ISL_529028, EPI_ISL_423312 and EPI_ISL_423313), GV (EPI_ISL_649919) and GH (EPI_ISL_422008 and EPI_ISL_728375) from the US that all do not contain N501Y are included for reference.
