## Supplementary Table 1 for "Distinct Patterns of Emergence of SARS-CoV-2 Spike Variants including N501Y in Clinical Samples in Columbus Ohio"

**Supplementary Table 1. Defining variants characteristics of SARS-CoV-2 clades, based on NextStrain terminology.**

| Gene | Nucleotide | Amino Acid | COH32* | B.1.1.7** | SA# |
|---|---|---|---|---|---|
| **G strain** | | | | | |
| S | A23403G | D614G | + | + | + |
| Upstream | C241T | NA | + | + | + |
| ORF1ab | C3037T | F924F | + | + | + |
| | C14408T | L4715L | + | + | + |
| **Clade 20B** | | | | | |
| N | 28881 GGG->AAC | RG203KR | - | + | - |
| **B.1.1.7 (20I/501Y.V1)** | | | | | |
| S | A23063T | N501Y | + | + | + |
| | 21765-21770del | HV69-70del | - | + | - |
| | 21991-21993del | Y144del | - | + | - |
| | C23271A | A570D | - | + | - |
| | C23604A | P681H | - | + | - |
| | C23709T | T716I | - | + | - |
| | T24506G | S982A | - | + | - |
| ORF8 | G28048T | R52I | + | + | - |
| | C27972T | Q27* | - | + | - |
| | A28111G | Y37C | - | + | - |
| N | 28280 GAT->CTA | D3L | - | + | - |
| | C28977T | S235F | - | + | - |
| **Clade 20C** | | | | | |
| ORF1ab | C1059T | T265I | + | - | + |
| ORF3a | G25563T | Q57H | + | - | + |
| **Clade 20G** | | | | | |
| ORF3a | G25907T | G172V | + | - | - |
| ORF8 | C27964T | S24L | + | - | - |
| N | C28472T | P67S | + | - | - |
| | C28869T | P199L | + | - | - |

*COH.20G/501Y, **20I/501Y.V1, #20H/501Y.V2
